# Highly pathogenic avian influenza H5 virus exposure in goats and sheep

**DOI:** 10.1101/2024.08.31.610397

**Authors:** Foong Ying Wong, Tahir Yaqub, Rong Zhang, Nadia Mukhtar, Hamda Pervaiz, Hafiz Usama Hussain Yawar, Mubashir Iqbal, Hassaan bin Aslam, Muhammad Waqar Aziz, Maham Akram, Sumbal Raza, Jenny G Low, Peter Cronin, Eric D Laing, Dolyce HW Low, Richard J Webby, Yvonne CF Su, Gavin JD Smith

## Abstract

The recent outbreaks of highly pathogenic avian influenza A(H5N1) virus in North and South America, including widespread infection of cattle in the United States, calls for an urgent assessment of the host range of influenza A viruses, particularly for subtypes of pandemic concern. We conducted a serological survey for binding antibodies to influenza A and B viruses in goats (n=452) and sheep (n=329) in Pakistan and found high seropositive rates for the hemagglutinin (HA) of avian influenza A viruses (AIV) H5 (23.9–34.0%), H7 (13.9– 37.1%), and H9 (17.0–34.7%). In contrast, there were low levels of seropositivity against the HA of human and swine pandemic H1N1/pdm09 (0.9–1.8%) in goats and against swine H3 (0.6%) in sheep. Notably, we observed high reactivity to the neuraminidase of human H1N1/2009 (57.8–60.6%) and swine H3N2 (14.0–14.4%), likely due to cross-reactivity with the N1 and N2 proteins of H5N1 and H9N2 AIVs, respectively. Interestingly, we also detected seropositivity against influenza B HA in both goats (7.1%) and sheep (4.6%). The presence of AIV antibodies in goats and sheep suggest these species represent previously unrecognized hosts for viruses of pandemic concern, revealing extensive gaps in our current understanding of the ecology of influenza A and B viruses.

## Introduction

Since March 2024, the outbreak of highly pathogenic avian influenza (HPAI) A(H5N1) virus in the United States has led to an unprecedented spread of infections in dairy cattle across multiple states^1-3^. This HPAI H5N1 is a new reassortant virus belonging to clade 2.3.4.4b, comprising gene segments from both Eurasian and North American progenitor viruses^4^. In contrast to most avian influenza A viruses that typically cause lower and upper respiratory infections in birds and animals, this HPAI H5N1 replicates efficiently in the epithelial cells of mammary glands of infected cows^3^. An experimental study showed that bovine HPAI H5N1 viruses induce severe weight loss and mortality in infected mice, but demonstrated inefficient transmission between ferrets without seroconversion in infected animals^5^. These studies raise concerns about the host species range of influenza A viruses, particularly those subtypes that pose zoonotic and pandemic risks^6^.

Following the emergence of the HPAI A/goose/Guangdong/1/1996 (H5N1) lineage in China in 1996, the virus has undergone rapid lineage diversification and extensive reassortment, evolving into numerous genotypes, causing multiple outbreaks in wild birds and domestic poultry across Asia, the Middle East and Europe^7-9^. Additionally, other major avian subtypes such as H7N9 and H9N2 have been circulating endemically, leading to sporadic outbreaks among poultry in mainland China and Southeast Asia^10-14^. Avian influenza A viruses are classified into low pathogenic (LPAI) and highly pathogenic strains, which are distinguished by the presence of polybasic cleavage sites in the hemagglutinin (HA) proteins. To date, only H5 and H7 subtype viruses are known to have HPAI variants. While LPAI viruses typically do not cause obvious clinical diseases in birds, HPAI H5N1 and H7N9 result in high mortality rates in poultry^11, 15, 16^.

Active serological surveillance and monitoring of avian influenza viruses in animals depends on screening for antibodies to influenza A viruses. Gold standard assays, such as enzyme-linked immunosorbent (ELISA), hemagglutination inhibition (HI) and microneutralization (MN) assays, are widely used to detect the presence of antibodies against influenza A viruses. However, these assays are labor-intensive, time-consuming, and the HI and MN assays typically require the use of live viruses, necessitating the use of a high containment facility for handling HPAI strains. The emergence of SARS-CoV-2 virus has spurred the rapid development of more efficient platforms for serological assessment. Multiplex serological assays have proven effective in evaluating neutralizing antibodies in humans against SARS-CoV-2 and other sarbecoviruses as well as in assessing antibody response following COVID-19 vaccinations or natural infections^17, 18^. We and others have previously developed antigen-based multiplex serological assays for the detection of antibodies against filovirus, henipavirus, and influenza viruses^19-21^.

Avian influenza A viruses circulate globally in wild and domestic birds and direct zoonotic transmission into humans is uncommon. However, seroprevalence of human antibodies to avian influenza A viruses (AIV) are consistently detected among poultry workers in many countries and regions including China, Egypt, Hong Kong, Pakistan, and Thailand^22-27^. Most serological studies have concentrated on commercial poultry and humans and there is a global lack of testing in domestic animals, and limited information available on AIV seroprevalence in other livestock, including bovid species such as goats and sheep^28, 29^.

Here, we collected over 700 sera samples from goats and sheep in Pakistan in 2023. A multiplex serological platform was employed to screen and evaluate sera samples against a panel of influenza HA and neuraminidase (NA) proteins. Our findings reveal that the sera from goats and sheep exhibited strong reactivity to avian influenza HA proteins, including HPAI H5 and LPAI H7 and H9, suggesting previous infections in these animals. In contrast, there was remarkably low level of reactivity to proteins of human H1N1/pdm09 and seasonal H3 viruses. These results indicate that both goat and sheep may be susceptible to avian influenza viruses, raising zoonotic concerns about their ecological role as hosts for avian influenza virus transmission globally.

## Results

Farms in four districts (Gujranwala, Kasur, Lahore, and Sheikhupura) in Punjab Province, Pakistan were selected as study sites (Fig. 1). A total of 781 serum samples were collected from goats (n=452) and sheep (n=329) from May to October 2023. These goats and sheep are primarily bred for meat production and typically slaughtered at 1.5–2 years of age. Serum samples were screened using a multiplex microsphere immunoassay comprised of nine hemagglutinin (HA) and three neuraminidase (NA) antigens of avian (H5, H7 and H9), human (H1N1/pdm09, seasonal H3, and influenza B), and swine influenza subtypes (Suppl. Table 1).

**Figure 1.**
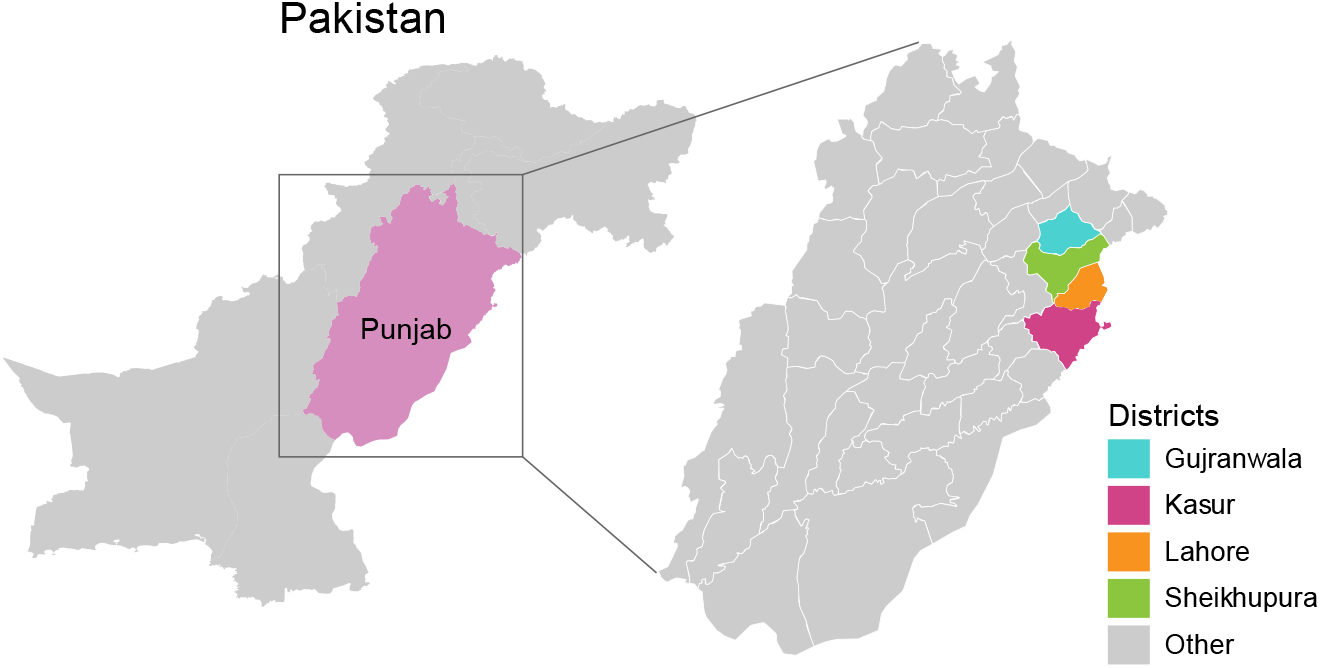
Geographical map of Punjab, Pakistan. Coloured areas indicate the location of Gujranwala, Kasur, Lahore and Sheikhupura districts from where goats and sheep were sampled.

High levels of seropositivity to AIV HA proteins were observed in both goat and sheep samples. Among the 452 goat serum samples collected, 124 samples (27.4%) tested positive for HPAI H5 clade 2.3.2.1c, 108 (23.9%) for H5 clade 2.3.4.4b, 63 (13.9%) for H7, and 77 samples (17.0%) for H9 (Fig. 2a, Suppl. Table 2). In contrast, there was little reactivity against influenza A viruses that circulate in humans and swine. All goat sera were negative against human and swine H3-HA, and only 8 (1.8%) and 4 (0.9%) of samples, respectively, were reactive to the human and swine H1N1/pdm09-HA antigen. However, 32 (7.1%) goat sera were seropositive to the HA protein of human B/Victoria virus. Similarly, among the 329 sheep serum screened, we detected 112 (34.0%) positive for antibodies to the HAs of H5 clade 2.3.2.1c, and 102 (31.0%) for H5 clade 2.3.4.4b (Fig. 2b, Suppl. Table 3). Additionally, 122 samples (37.1%) were positive against H7-HA and 114 (34.7%) against H9-HA antigens. No sheep sera were positive against the HAs of human and swine H1N1/pdm09 or human H3. Only 2 (0.6%) sheep serum samples were positive against the swine H3-HA, while 15 (4.6%) were positive for the B/Victoria HA protein.

**Figure 2.**
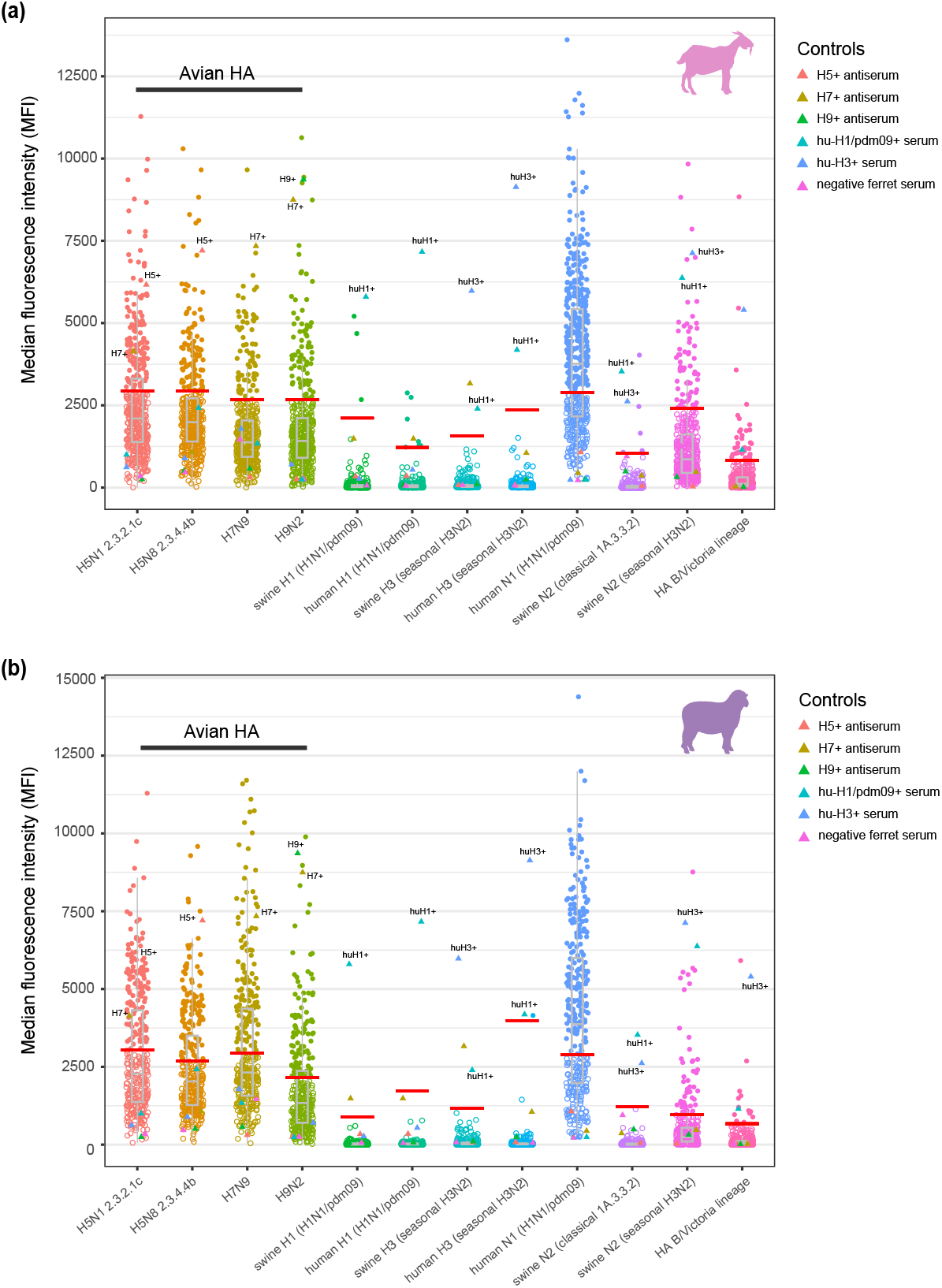
Serological dynamics of goat and sheep sera against a panel of influenza virus antigens. Median fluorescence intensity (MFI) values were measured for individual **(a)** goat (n=452) and **(b)** sheep (n=329) serum samples using a multiplex microsphere immunoassay. For each sample, the reactivity of sera was assessed against 12 influenza A and B virus antigens including nine HA proteins from HPAI A(H5N1) clade 2.3.2.1c, HPAI A(H5N8) clade 2.3.4.4b, A(H7N9), A(H9N2), swine H1N1/pdm09, human H1N1/pdm09, swine H3N2), human seasonal H3N2, and human B/Victoria viruses, and three NA proteins from human H1N1/pdm09, classical swine H1N2 and swine H3N2 viruses. Solid colored circles represent seropositive samples, while open circled dots denote seronegative samples. Red lines indicate the cutoff value for each antigen determined using an expectation–maximization algorithm (see Suppl Fig. 1 and Suppl. Table 1). Box-plots indicate the median and quartiles of the MFI distribution for each antigen. Positive and negative serum controls are denoted by colored triangles, and included 3 ferret antisera for H5, H7 and H9, 1 PCR-positive H1N1/pdm09 patient, 1 PCR-positive H3N2 patient and 1 negative ferret control.

For both goat and sheep sera there was high positivity to the NAs of human H1N1/pdm09 and the swine seasonal H3N2 viruses (Fig. 2, Suppl. Tables 2 and 3). A significant proportion of goat (274/452, 60.6%,) and sheep (190/329, 57.8%) displayed reactivity to the H1N1/pdm09-NA antigen, while there was lower reactivity to the NA of swine seasonal H3N2 in both goats and sheep (14.0–14.4%), and minimal (1.3%) positives to the classical (clade 1A.3.3.2) swine N2-NA. This pattern of seropositivity against the NA antigens in our assay is possibly due to cross-reactivity with the N1-NA of H5N1 viruses and the N2-NA of H9N2 viruses that are endemic in birds in the region. The positive controls had reactivity patterns as expected, although the H7 ferret antiserum showed broad reactivity with multiple, but not all, antigens (Fig. 2). The positive controls for the H1, H3, N1 and N2 proteins were sera collected from patients 21 days after they were confirmed to have either acute H1N1/pdm09 or H3N2 virus infection. As humans are likely to have had multiple influenza A virus infections throughout their lifetime, it is unsurprising that some cross-reactivity was observed in these positive controls, such as the human post H3 infection control sera showing positivity against human H1-HA.

Next, we evaluated and compared the patterns of seropositivity among HA and NA antigens included in our multiplex assay. For sera that were positive to a specific influenza antigen, we calculated their reactivity towards other antigens (Fig. 3). For example, in goats there were 124 sera positive against the HA of H5N1 clade 2.3.2.1c, and of these 90 were also positive against clade 2.3.2.4b, while there were 43 and 62 positives against the H7 and H9 antigens, respectively (Fig. 3a). In both goats and sheep, the majority of sera positive to the HA of avian H5N1 clade 2.3.2.1c strain were also positive for the HA of H5N8 clade 2.3.4.4b, and vice versa (Fig. 3). A substantial proportion of H5-HA positive samples were also positive for H9-HA. Notably, the H7 positive sera in both goats and sheep showed lower cross-positivity with H5-HA and H9-HA. These patterns may reflect the infection history of these animals, where they have been exposed to all three AIV subtypes or it could be due to cross-reaction of the AIV antigens which include the more conserved HA2 domain of the influenza A HA protein. There were no or very few samples that were positive against AIV HA that were also positive to mammalian H1N1/pdm09-HA and H3-HA from humans and swine (Suppl. Fig. 1).

We show a substantial number of sera positive against the NA of H1N1/pdm09 in both goats and sheep that were also positive against the H5, H7 and H9 antigens (Fig. 3e and k). Of the 274 goat and 192 sheep sera positive against H1N1/pdm09 N1-NA, 124 and 112 sera, respectively, were also seropositive for the HA of H5N1 clade 2.3.2.1c. Given that only 12 goat sera and no sheep sera were positive against H1N1/pdm09 H1-HA, this shared positivity is likely due to cross-reactivity of the H1N1/pdm09 N1-NA with the N1-NA of H5N1 clade 2.3.2.1c viruses. In addition, H1N1/pdm09 N1-NA positive sera were also reactive against the HA proteins of H5N8 (goats: 98; sheep: 100), H7N9 (goats: 53; sheep: 96) and H9N2 (goats: 67; sheep: 94) viruses. A similar pattern was observed for the N2-NA antigens, which had positive samples that were also positive for the AIV HAs and H1N1/2009 N1-NA (Fig. 3). The cross-positivity of sera against N1– and N2-NAs with H5-, H7– and H9-HAs supports the contention that we are observing an infection history comprised of multiple AIV subtype exposures.

**Figure 3.**
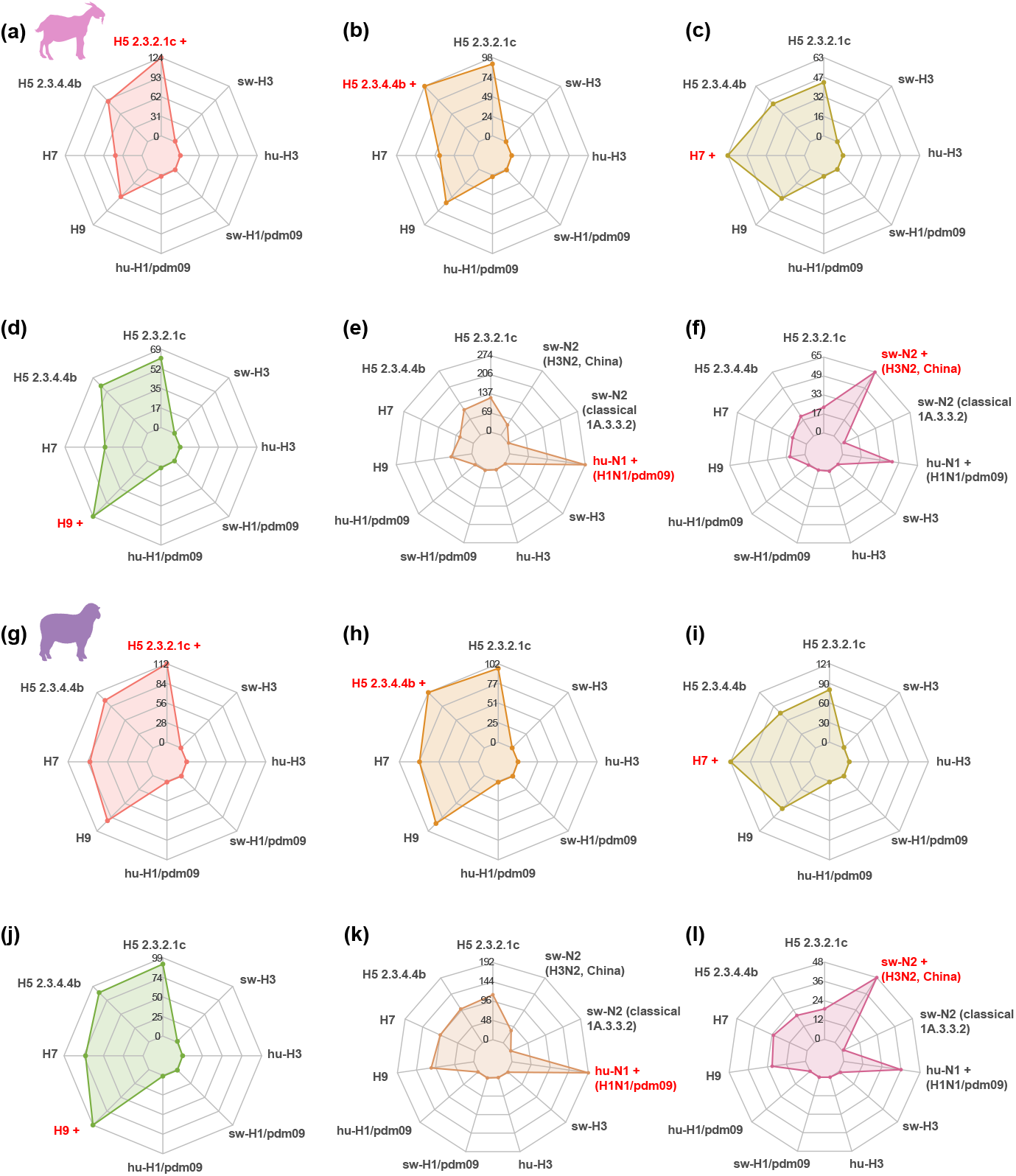
Comparison of patterns of seropositivity of HA and NA antigens in goat and sheep. The total number of positive serum samples from **(a-f)** goat and **(g-l)** sheep that also tested positive for another antigen are shown. In each radar plot, the red font with a plus sign indicates the antigen with positive samples that are being compared for seropositivity to the other antigens, shown in black font. The numbers in the centre of each plot represent the number of seropositive samples for each antigen. Abbreviations: hu-H1/pdm09, human H1N1/pdm09; sw-H1/pdm09, swine H1N1/pdm09; hu-H3, human seasonal H3N2; sw-H3, swine H3N2, sw-N2, swine H3N2 and sw-N2, classical swine H1N2.

We further compared the prevalence among districts in Pakistan. Both goat and sheep from all four districts consistently showed higher seropositivity rates for avian HA subtypes (H5, H7 and H9) compared to the human H1N1/pdm09 and seasonal H3 (Fig. 4a and b). Among these districts, Sheikhupura showed the highest seropositivity rates for anti-H5 and anti-H7 antibodies in goats and sheep. Additionally, the level of anti-N1 antibodies remained remarkably high in both goats (55.4–65.1%) and sheep (41.4–75.5%) across all four districts, compared to anti-N2 antibodies (goats: 4.1–12.8% and sheep: 8.6–21.6%) (Fig. 4a and b; Suppl. Tables 2 and 3). We also observed that the range of median fluorescence intensity (MFI) values for seropositive samples were relatively consistent among different antigens in our assay (Fig. 4c). However, there were significant differences in the MFI values between goat and sheep positive sera, when a comparison was possible, except for the HA of H5N8 and influenza B which showed no significant difference (Fig. 4c). For the HA of H5N1 2.3.2.1c and H7N9, and the NA of H1N1/pdm09, sheep had higher MFIs compared to goats, suggesting that sheep may have had more recent exposure to H5N1 and H7 viruses. The opposite was observed for the HA of H9N2 and the NA of swine seasonal H3N2, suggesting that sampled goats may have been recently infected with H9N2 viruses.

**Figure 4.**
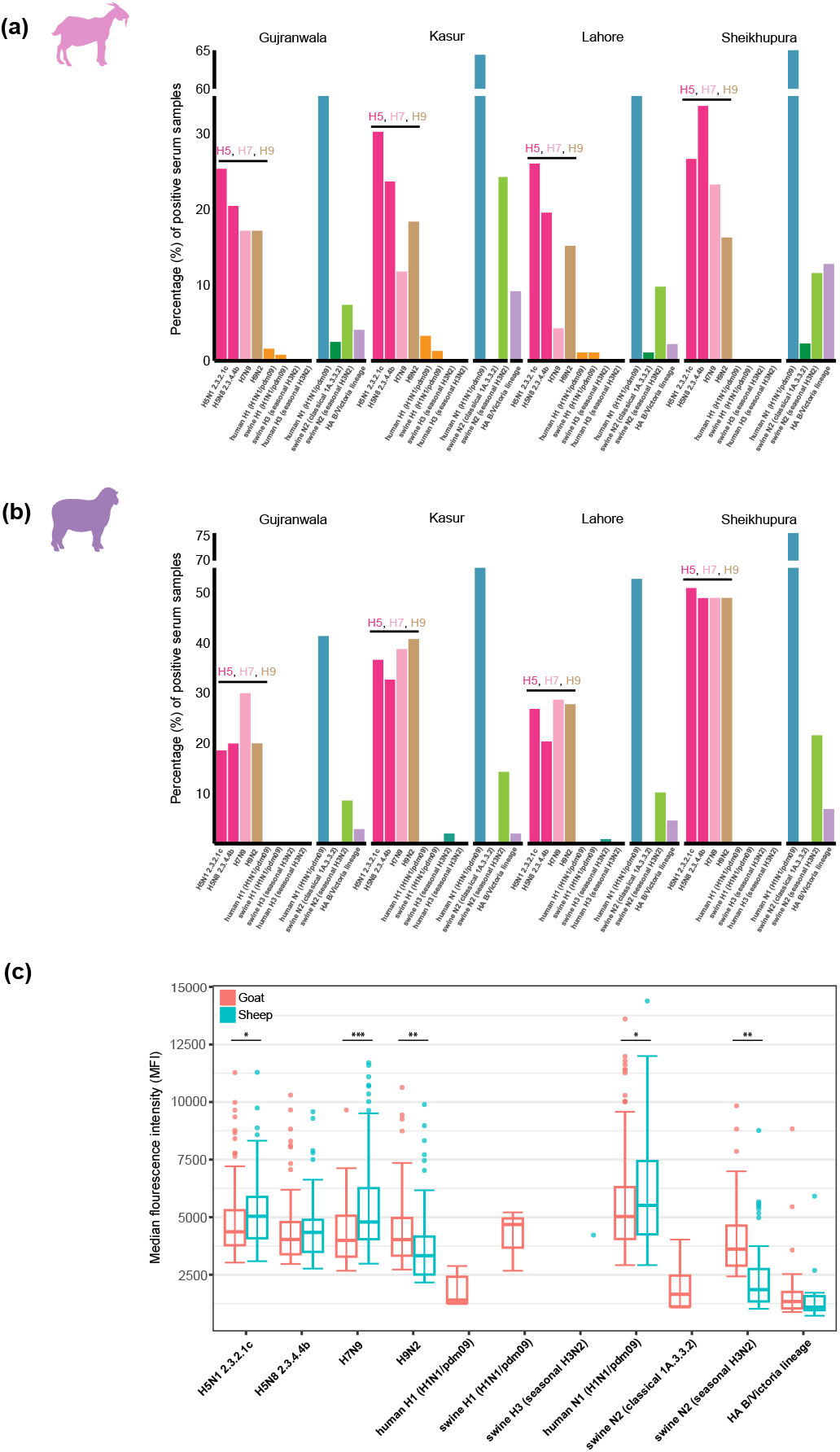
Distribution of seroprevalence among geographical locations and host species. The seropositivity of influenza A and B viruses is shown as percentage in **(a)** goat and **(b)** sheep across the four districts in Punjab, Pakistan. Coloured vertical bars represent different influenza virus antigens corresponding to specific subtypes or lineages. **(c)** Boxplot showing distribution of median fluorescence intensity (MFI) values of positive samples for goat and sheep. The central line within each box represents the median value, while the upper and lower edges of the box correspond to the third and first quartiles, respectively, whiskers indicate data points within 1.5 times the interquartile range from the quartiles, with data points outside of this range shown as individual dots. Red coloured box plots represent goat, while blue-coloured box plots represent sheep. Statistical significance between MFI distributions of goat and sheep was assessed using the Kruskal-Wallis with Dunns post-hoc test for multiple comparisons. Significance levels are indicated by asterisks: * P ≤ 0.05, ** P ≤ 0.001, and *** P ≤ 0.0001.

## Discussion

Here we detect a high degree of AIV seroprevalence in goats and sheep from Pakistan, suggesting these animals have been exposed to past H5, H7, and H9 infection. Disentangling evidence of multiple infections from cross-reactive antibodies generated after a single infection, which are expected with an antibody binding assay such as we used, is difficult. The breadth of antibody specificity in post-infection goat and sheep antisera is unknown. Interestingly, these goats and sheep samples were collected in areas with commercial chicken farms, that are known to support the endemic circulation of H5, H7 and H9 AIV in domestic poultry such as chicken and quail in Pakistan^30-33^. Pakistan has intensive production of goats and sheep, with respectively over 80 and 31 million animals raised annually^34^. Since goats and sheep are not vaccinated against influenza A virus in Pakistan, their exposure to AIV highlights the potential risk of virus transmission from chickens where goat and sheep farming are often located near chicken rearing units. Seasonal migration of small ruminants, in particular sheep, is not uncommon by nomadic movement across the Afghanistan and Pakistan border^35^, highlighting the need for urgent influenza surveillance in countries with significant populations of goats, sheep and other bovids.

Global production of goats (1.1 billion head in 2021) and sheep (1.3 billion) are concentrated in Asia and Africa, 95% and 77%, respectively^36^, with China and India the top two largest producers for both species. Given the recent emergence of H5N1 infection in cattle in the USA, the detection of anti-AIV antibodies in goats and sheep is of major public health concern. The failure to recognize the potential role of a broader range of livestock species as potential hosts for AIV is a major blind spot in pandemic preparedness planning. Currently, there is an almost complete lack of knowledge of the presence and distribution of influenza virus receptors, a major determinant of susceptibility to influenza infection, in many livestock species. It was only after the emergence of H5N1 in cattle that detailed investigation of receptor distribution has occurred, which revealed the presence of alpha 2,3 and alpha 2,6 in mammary glands^37, 38^. These recent discoveries suggest that if the outbreak in cattle is not controlled, that cattle could act as an ideal intermediate host for adaptation of AIV to mammalian receptors.

Potential susceptibility of sheep to influenza A viruses has been previously demonstrated through the productive replication of influenza A/H10N7 virus on sheep respiratory tissues that resulted in extensive cytopathic effect during virus culture^39^. Furthermore, antibodies against influenza A virus have been detected in goat milk and blood when mammary glands were injected with influenza A/PR/8/34 virus^40^. Additionally, goats, sheep and camels have shown seropositivity to influenza D virus in Ethiopia and USA^41, 42^. Seroconversion to influenza H9 virus has also been observed in water buffalo (*Bubalus bubalis*) in Iran, with 14 of 80 (17.5%) slaughtered animals positive for anti-H9 antibodies in a hemagglutinin inhibition assay^43^.

Our results emphasize the importance of expanded surveillance and disease monitoring in a wider variety of hosts, particularly livestock, in areas where there is endemic circulation of AIV of pandemic concern. Experimental studies are also needed to investigate receptor distributions within these animals to better inform their potential role in virus adaptation to receptors prevalent in the human respiratory tract. Until such studies are conducted, our incomplete understanding of the ecology of influenza A viruses will remain a major impediment to any pandemic preparedness efforts.

## Methods

### Sample collection

Four sites were surveyed for the sample collection in Punjab Province, Pakistan. A total of 794 serum samples were collected from goats (n=459) and sheep (n=335) from May–October 2023 that were co-housed or residing near the vicinity of commercial chicken farms. Approximately 5 ml of blood was drawn with a sterile syringe from the jugular vein of goats and sheep and collected in a serum separation vacutainer. Serum was separated from blood samples and stored at –20°C until further processing. All procedures were approved by the Ethical Review Committee at the University of Veterinary and Animal Sciences Lahore in Pakistan (DR/361).

### Multiplex serological assay

To assess the prevalence of antibodies against a panel of avian, human and swine influenza A, and influenza B viruses, individual samples were tested via a multiplex microsphere immunoassay using the Luminex MAGPIX platform that was adapted from previously established assays^19, 20^. The influenza virus HA and NA proteins were either commercially available (Sino Biological Inc) or synthesized by GenScript Biotech (Suppl. Table 1), and altered to include polyhistidine tags and to remove transmembrane regions. Briefly, 30μg of recombinant proteins were coupled on Bio-Plex Pro™ carboxylated microsphere beads (Bio-Rad) at 24 μg millions^-1^ beads, for 2 h at room temperature in the dark with agitation. These coupled beads were then incubated with sera samples at a dilution of 1:100 for 45 min at room temperature with agitation. After incubation, the samples were washed and incubated with biotinylated-Protein A and biotinylated-Protein G (Thermo Fisher Scientific), followed by streptavidin-phycoerythrin (PE) (Bio-Rad). After three washes, the final median fluorescence intensity (MFI) values were measured using the MAGPIX system (Luminex) with xPONENT 4.3 software, according to the manufacturer’s instructions.

We also selected representative ferret and human antiserum samples as controls (Suppl. Table 4). For human antisera controls, respiratory swabs and blood samples were collected from volunteers with respiratory symptoms at their first visit and followed up at convalescence on Day 21±5 (CIRB Ref: 2018/2425). Influenza positive samples were confirmed by PCR. Animal antisera for different avian influenza H5, H7, and H9 subtypes, was generated by experimentally infecting ferrets with known AIV subtypes.

### Data analysis

Antigen-antibody complexes were qualitatively measured as median fluorescence intensities (MFI) using the MAGPIX system (Luminex) with xPONENT 4.3 software following the manufacturer’s instructions. Cutoffs to identify positive samples were determined using an expectation–maximization algorithm applying the Mclust function in R package mclust^44^. Prior to classification, serum samples with an MFI >8,000 to more than one protein were removed, resulting in a final sample set of 781 sera (goat, n=452 and sheep, n=329). For each protein, the MFI values of all samples were classified into a maximum of 4 clusters, after which the log-likelihood corresponding to the specific number of clusters was obtained and the optimal model selected according to Bayesian Information Criteria (Suppl. Figs 2 and 3; Suppl. Table 4). The samples classified into the last group were considered positive. To investigate patterns of seropositivity among HA and NA antigens, we took the samples that were positive to a specific influenza antigen and plotted the number of those that were also positive for other antigens. The R package fmsb was used to generate radar plots while all other figures were produced using ggplot2. Statistical significance was tested using Kruskal-Wallis with Dunns post-hoc test in the PMCMRplus package also in R.

## Acknowledgements

This study was supported by the Duke-NUS Signature Research Programme funded by Ministry of Health, Singapore, Singhealth Duke-NUS Global Health Institute research grant RGA(Khoo)/2022/0010, and by contract 75N93021C00016 from the National Institute of Allergy and Infectious Diseases, National Institutes of Health, Department of Health and Human Services, USA. We acknowledge the Livestock and Dairy Development Department of Punjab, Pakistan, and the dairy farmers for their invaluable assistance in the collection of ruminant samples for this project. We thank Dr Michael Zeller at Duke-NUS for providing the initial scripts for data analyses.

## Author contributions

T.Y., Y.C.F.S and G.J.D.S. conceived the study; N.D., H.P., H.U.H.Y., M.I., H.A., M.W.A, M.A and S.R. designed field work and collected samples; J.G.L. provided samples; F.Y.W. and D.H.W.L. performed experiments; F.Y.W., R.Z., P.C., Y.C.F.S and G.J.D.S. analysed data; F.Y.W., Y.C.F.S. and G.J.D.S. wrote the draft, with input from E.D.L. and R.J.W. All authors commented on and reviewed the manuscript.

## Notes

### Competing Interest Statement

The authors have declared no competing interest.

## References

1. Burrough, E.R. et al. Highly Pathogenic Avian Influenza A(H5N1) Clade 2.3.4.4b Virus Infection in Domestic Dairy Cattle and Cats, United States, 2024. Emerg Infect Dis 30, 1335–1343 (2024).

2. Nguyen, T.-Q. et al. Emergence and interstate spread of highly pathogenic avian influenza A(H5N1) in dairy cattle. bioRxiv, 2024.2005.2001.591751 (2024).

3. Caserta, L.C. et al. Spillover of highly pathogenic avian influenza H5N1 virus to dairy cattle. Nature (2024).

4. Worobey, M. et al. in Preliminary report on genomic epidemiology of the 2024 H5N1 influenza A virus outbreak in U.S. cattle (Part 1 of 2) (virological.org, 2024) (2024).

5. Eisfeld, A.J. et al. Pathogenicity and transmissibility of bovine H5N1 influenza virus. Nature (2024).

6. James, J., Sealy, J.E. & Iqbal, M. A Global Perspective on H9N2 Avian Influenza Virus. Viruses 11, 620 (2019).

7. Global Consortium for, H.N. & Related Influenza, V. Role for migratory wild birds in the global spread of avian influenza H5N8. Science 354, 213–217 (2016).

8. Li, Y.T., Su, Y.C.F. & Smith, G.J.D. H5Nx Viruses Emerged during the Suppression of H5N1 Virus Populations in Poultry. Microbiol Spectr, e0130921 (2021).

9. Xie, R. et al. The episodic resurgence of highly pathogenic avian influenza H5 virus. Nature 622, 810–817 (2023).

10. Yang, L. et al. Genesis and Spread of Newly Emerged Highly Pathogenic H7N9 Avian Viruses in Mainland China. J Virol 91 (2017).

11. Lam, T.T. et al. Dissemination, divergence and establishment of H7N9 influenza viruses in China. Nature 522, 102–105 (2015).

12. Choi, Y.K. et al. Continuing evolution of H9N2 influenza viruses in Southeastern China. J Virol 78, 8609–8614 (2004).

13. Pu, J. et al. Evolution of the H9N2 influenza genotype that facilitated the genesis of the novel H7N9 virus. Proc Natl Acad Sci U S A 112, 548–553 (2015).

14. Millman, A.J. et al. Detecting Spread of Avian Influenza A(H7N9) Virus Beyond China. Emerg Infect Dis 21, 741–749 (2015).

15. Isoda, N. et al. Re-Invasion of H5N8 High Pathogenicity Avian Influenza Virus Clade 2.3.4.4b in Hokkaido, Japan, 2020. Viruses 12 (2020).

16. Kim, H.R. et al. Highly pathogenic avian influenza (H5N1) outbreaks in wild birds and poultry, South Korea. Emerg Infect Dis 18, 480–483 (2012).

17. Tan, C.W. et al. SARS-CoV-2 Omicron variant emerged under immune selection. Nat Microbiol 7, 1756–1761 (2022).

18. Chia, W.N. et al. Dynamics of SARS-CoV-2 neutralising antibody responses and duration of immunity: a longitudinal study. Lancet Microbe 2, e240–e249 (2021).

19. Dovih, P. et al. Filovirus-reactive antibodies in humans and bats in Northeast India imply zoonotic spillover. PLOS Neglected Tropical Diseases 13, e0007733 (2019).

20. Laing, E.D. et al. Serologic Evidence of Fruit Bat Exposure to Filoviruses, Singapore, 2011-2016. Emerg Infect Dis 24, 114–117 (2018).

21. Germeraad, E. et al. The development of a multiplex serological assay for avian influenza based on Luminex technology. Methods 158, 54–60 (2019).

22. Jiang, L. et al. Cross-reactive antibodies against H7N9 and H5N1 avian influenza viruses in Thai population. Asian Pac J Allergy Immunol 35, 20–26 (2017).

23. Yang, P. et al. Avian influenza A(H7N9) and (H5N1) infections among poultry and swine workers and the general population in Beijing, China, 2013–2015. Scientific Reports 6, 33877 (2016).

24. Gomaa, M.R. et al. Serological Evidence of Human Infection with Avian Influenza A H7virus in Egyptian Poultry Growers. PLoS One 11, e0155294 (2016).

25. Bridges, C.B. et al. Risk of Influenza A (H5N1) Infection among Poultry Workers, Hong Kong, 1997–1998. The Journal of Infectious Diseases 185, 1005–1010 (2002).

26. Chen, X. et al. Serological evidence of human infections with highly pathogenic avian influenza A(H5N1) virus: a systematic review and meta-analysis. BMC Medicine 18, 377 (2020).

27. Tahir, M.F. et al. Seroprevalence and risk factors of avian influenza H9 virus among poultry professionals in Rawalpindi, Pakistan. J Infect Public Health 13, 414–417 (2020).

28. Gupta, S.D., Fournié, G., Hoque, M.A. & Henning, J. Farm-Level Risk Factors Associated With Avian Influenza A (H5) and A (H9) Flock-Level Seroprevalence on Commercial Broiler and Layer Chicken Farms in Bangladesh. Front Vet Sci 9, 893721 (2022).

29. Lin, T.N. et al. Serological evidence of avian influenza virus subtype H5 and H9 in live bird market, Myanmar. Comp Immunol Microbiol Infect Dis 73, 101562 (2020).

30. Ali, M. et al. Genetic Characterization of Highly Pathogenic Avian Influenza A(H5N8) Virus in Pakistani Live Bird Markets Reveals Rapid Diversification of Clade 2.3.4.4b Viruses. Viruses 13 (2021).

31. Ali, M. et al. Avian Influenza A(H9N2) Virus in Poultry Worker, Pakistan, 2015. Emerging infectious diseases 25, 136–139 (2019).

32. Ali, M. et al. Prevalence and Phylogenetics of H9N2 in Backyard and Commercial Poultry in Pakistan. Avian Diseases 62, 416–424 (2018).

33. Channa, A.A., Tariq, M., Nizamani, Z.A. & Kalhoro, N.H. Prevalence of avian influenza H5, H7, and H9 viruses in commercial layers in Karachi, Pakistan. Iran J Vet Res 22, 352–355 (2021).

34. Mustafa, M.Y. & Ali, S.S. Prevalence of infectious diseases in local and fayoumi breeds of rural poultry (Gallus domesticus). Punjab Univ. J. Zool 20, 177–180 (2005).

35. Wajid, A., Chaudhry, M., Rashid, H.B., Gill, S.S. & Halim, S.R. Outbreak investigation of foot and mouth disease in Nangarhar province of war-torn Afghanistan, 2014. Sci Rep 10, 13800 (2020).

36. FAO, U.N. in FAOSTAT: Crops and livestock products (2022), https://www.fao.org/faostat (accessed 08 August 2024).

37. Nelli, R.K. et al. Sialic Acid Receptor Specificity in Mammary Gland of Dairy Cattle Infected with Highly Pathogenic Avian Influenza A(H5N1) Virus. Emerg Infect Dis 30, 1361–1373 (2024).

38. Chopra, P. et al. Receptor Binding Specificity of a Bovine A(H5N1) Influenza Virus. bioRxiv (2024).

39. Mazzetto, E. et al. Replication of Influenza D Viruses of Bovine and Swine Origin in Ovine Respiratory Explants and Their Attachment to the Respiratory Tract of Bovine, Sheep, Goat, Horse, and Swine. Front Microbiol 11, 1136 (2020).

40. Mitchell, C.A., Walker, R.V. & Bannister, G.L. Studies relating to the formation of neutralizing antibody following the propagation of influenza and Newcastle disease virus in the bovine mammary gland. Can J Microbiol 2, 322–328 (1956).

41. Quast, M. et al. Serological evidence for the presence of influenza D virus in small ruminants. Vet Microbiol 180, 281–285 (2015).

42. Murakami, S. et al. Influenza D Virus Infection in Dromedary Camels, Ethiopia. Emerg Infect Dis 25, 1224–1226 (2019).

43. Tajik, J., Tavakoli, H. & Soltani, D. Serological Investigation of H9N2 Avian Influenza Virus in Slaughtered Water Buffaloes (Bubalus bubalis) in Khuzestan, Iran. Archives of Razi Institute 74, 77–82 (2019).

44. Scrucca, L., Fop, M., Murphy, T.B. & Raftery, A.E. mclust 5: Clustering, Classification and Density Estimation Using Gaussian Finite Mixture Models. R j 8, 289–317 (2016).

