## Supplementary material for "Highly pathogenic avian influenza H5 virus exposure in goats and sheep"

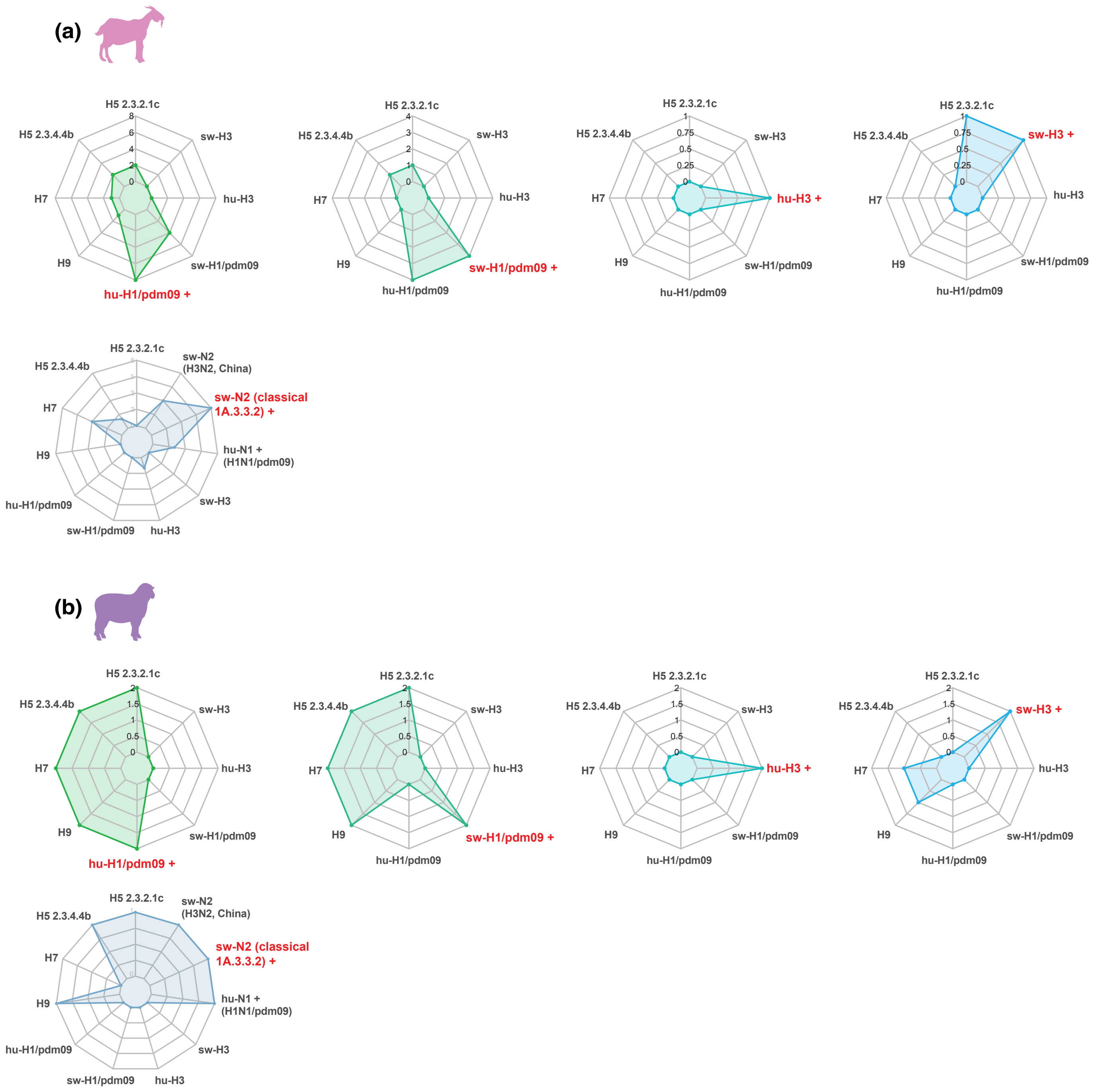


**Supplementary Figure 1.** Radar plots showing the patterns of seropositivity of HA and NA antigens. Total number of HA positive serum samples from **(a)** goat and **(b)** sheep that are positive for another antigen in our assay.

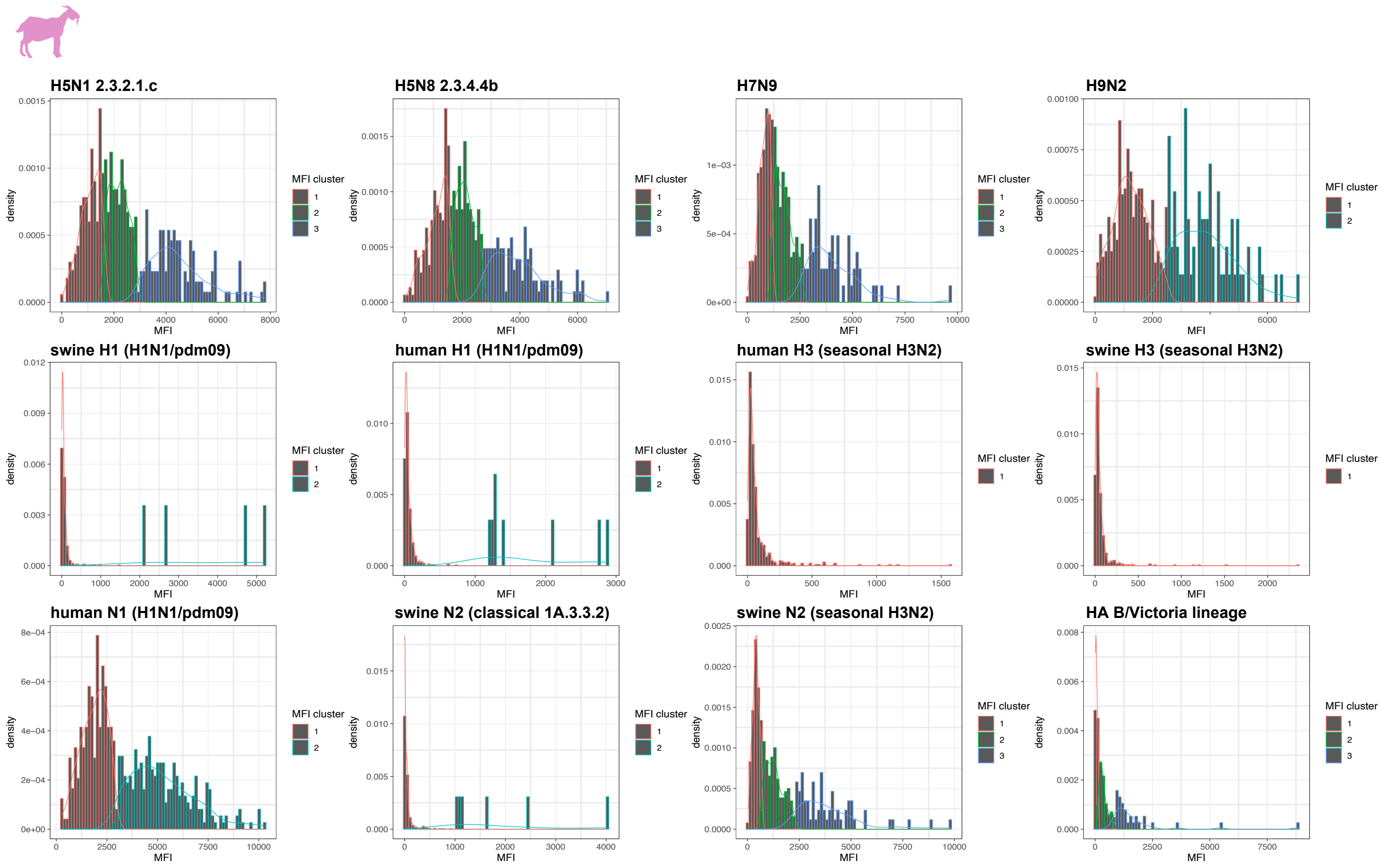


**Supplementary Figure 2.** Determination of median fluorescence intensity (MFI) cutoff values for goat sera against influenza A and B virus antigens.

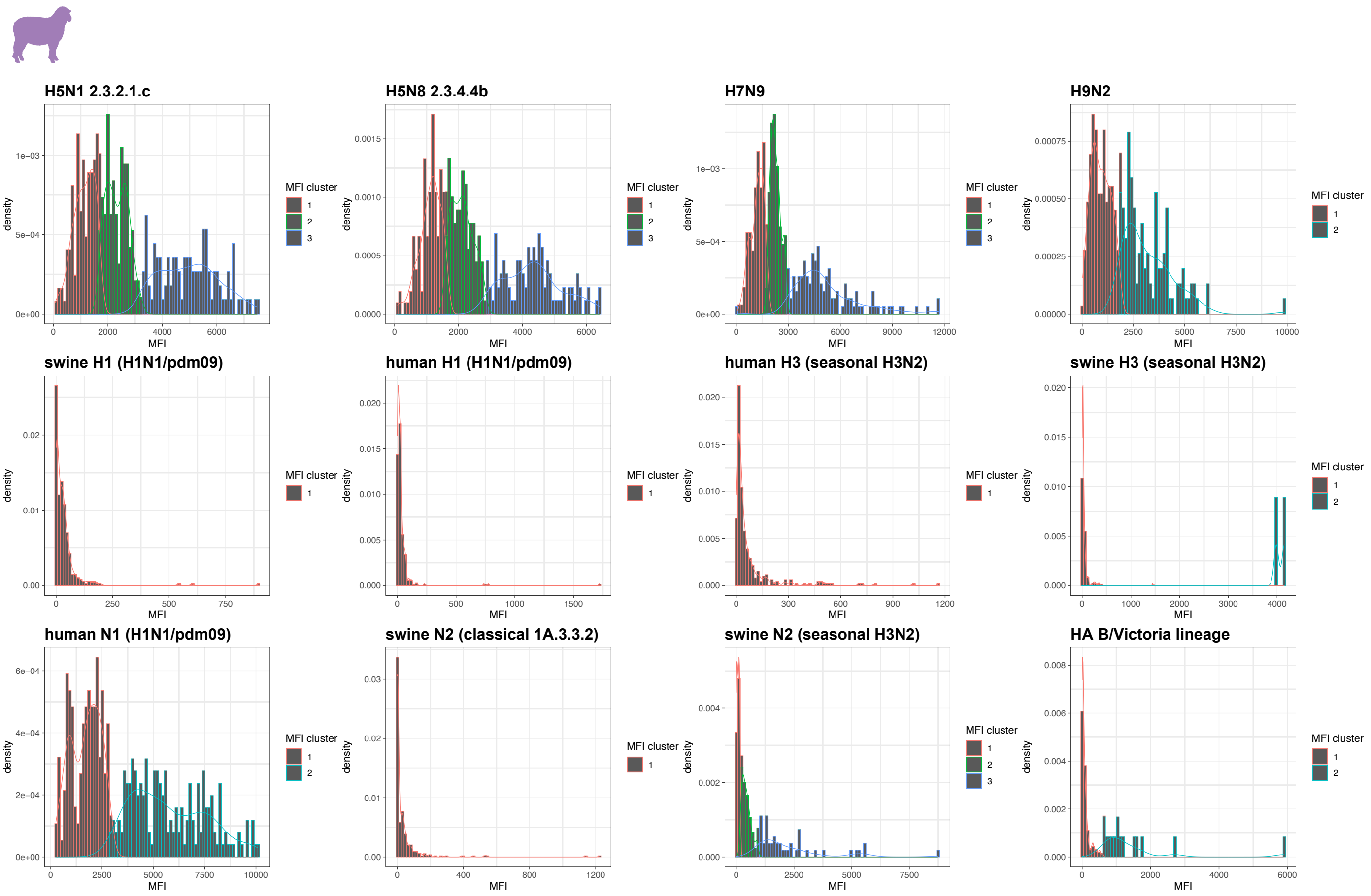


**Supplementary Figure 3.** Determination of median fluorescence intensity (MFI) cutoff values for sheep sera against influenza A and B virus antigens.

**Supplementary Table 1.** Influenza A and B virus antigens conjugated Bio-Plex beads used in multiplex assay.

| Recombinant protein | Subtype | Virus | GenBank accession no. | MFI cutoff value for each antigen* |
|---|---|---|---|---|
| H5-HA | H5N1 (2.3.2.1c) | A/Cambodia/NPH230032/2023 | EPI2419700 | 3046 |
| H5-HA | H5N8 (2.3.4.4b) | A/Astrakhan/3212/2020 | EPI1846961 | 2688 |
| H7-HA | H7N9 | A/Anhui/1/2013 | EPI439507 | 2944 |
| H9-HA | H9N2 (Y280) | A/Hong Kong/3239/2008 | ADC41863.1 | 2156 |
| H1-HA | H1N1/pdm09 | A/California/07/2009 | ACP41953.1 | 895 |
| H1-HA | H1N1/pdm09 | A/swine/Cambodia/PFC63/2021 | OQ460186 | 1721 |
| H3-HA | Human H3N2 (seasonal) | A/Cambodia/e0826360/2020 | EPI1837753 | 1163 |
| H3-HA | Swine H3N2 (seasonal) | A/swine/Thailand/CU-P53/2012 | AGG53108 | 3986 |
| N1-NA | H1N1/pdm09 | A/California/07/2009 | ACP41107.1 | 2882 |
| N2-NA | Swine H1N2 (classical 1A.3.3.2) | A/swine/Alberta/SD0217/2017 | QAX25989 | 1224 |
| N2-NA | Swine H3N2 (seasonal) | A/swine/China/JG20/2019 | QGZ99158 | 976 |
| B-HA | B/Victoria | B/Austria/1359417/2021 | EPI1868375 | 674 |

*Determined using an expectation–maximization algorithm

**Supplementary Table 2.** Seroprevalence of influenza A and B viruses in goat sera from Pakistan, May–October 2023.

| Goats | Gujranwala (*n*=122)<br>No. of positive (%) | Kasur (*n*=152)<br>No. of positive (%) | Lahore (*n*=92)<br>No. of positive (%) | Sheikhupura (*n*=86)<br>No. of positive (%) | Overall (*n*=452)<br>No. of positive (%) |
|---|---|---|---|---|---|
| **Hemagglutinin (HA)** | | | | | |
| A/Cambodia/NPH230032/2023 (H5N1 2.3.2.1c) | 31 (25.4%) | 46 (30.3%) | 24 (26.1%) | 23 (26.7%) | 124 (27.4%) |
| A/Astrakhan/3212/2020 (H5N1 2.3.4.4b) | 25 (20.5%) | 36 (23.7%) | 18 (19.6%) | 29 (33.7%) | 108 (23.9%) |
| A/Anhui/1/2013 (H7N9) | 21 (17.2%) | 18 (11.8%) | 4 (4.3%) | 20 (23.3%) | 63 (13.9%) |
| A/Hong Kong/3239/2008 (H9N2) | 21 (17.2%) | 28 (18.4%) | 14 (15.2%) | 14 (16.3%) | 77 (17.0%) |
| A/California/07/2009 (H1N1/pdm09) | 2 (1.6%) | 5 (3.3%) | 1 (1.1%) | - | 8 (1.8%) |
| A/swine/Cambodia/PFC63/2021 (H1N1/pdm09) | 1 (0.8%) | 2 (1.3%) | 1 (1.1%) | - | 4 (0.9%) |
| A/Cambodia/e0826360/2020 (seasonal H3N2) | - | - | - | - | - |
| A/swine/Thailand/CU-P53/2012 (seasonal H3N2) | - | - | - | - | - |
| **Neuraminidase (NA)** | | | | | |
| A/California/07/2009 (H1N1/pdm09) | 69 (56.6%) | 98 (64.5%) | 51 (55.4%) | 56 (65.1%) | 274 (60.6%) |
| A/swine/Alberta/SD0217/2017 (H1N2 classical 1A.3.3.2) | 3 (2.5%) | - | 1 (1.1%) | 2 (2.3%) | 6 (1.3%) |
| A/swine/China/JG20/2019 (seasonal H3N2) | 9 (7.4%) | 37 (24.3%) | 9 (9.8%) | 10 (11.6%) | 65 (14.4%) |
| **Influenza B hemagglutinin (HA)** | | | | | |
| B/Austria/1359417/2021 | 5 (4.1%) | 14 (9.2%) | 2 (2.2%) | 11 (12.8%) | 32 (7.1%) |

**Supplementary Table 3.** Seroprevalence of influenza A and B viruses in sheep sera from Pakistan, May–October 2023.

| Sheep | Gujranwala (*n*=70) | Kasur (*n*=49) | Lahore (*n*=108) | Sheikhupura (*n*=102) | Overall (*n*=329) |
|---|---|---|---|---|---|
| | No. of positive (%) | No. of positive (%) | No. of positive (%) | No. of positive (%) | No. of positive (%) |
| **Hemagglutinin (HA)** | | | | | |
| A/Cambodia/NPH230032/2023 (H5N1 2.3.2.1c) | 13 (18.6%) | 18 (36.7%) | 29 (26.9%) | 52 (51.0%) | 112 (34.0%) |
| A/Astrakhan/3212/2020 (H5N1 2.3.4.4b)) | 14 (20.0%) | 16 (32.7%) | 22 (20.4%) | 50 (49.0%) | 102 (31.0%) |
| A/Anhui/1/2013 (H7N9) | 21 (30.0%) | 19 (38.8%) | 31 (28.7%) | 50 (49.0%) | 122 (37.1%) |
| A/Hong Kong/3239/2008 (H9N2) | 14 (20.0%) | 20 (40.8%) | 30 (27.8%) | 50 (49.0%) | 114 (34.7%) |
| A/California/07/2009 (H1N1/pdm09) | - | - | - | - | - |
| A/swine/Cambodia/PFC63/2021 (H1N1/pdm09) | - | - | - | - | - |
| A/Cambodia/e0826360/2020 (seasonal H3N2) | - | - | - | - | - |
| A/swine/Thailand/CU-P53/2012 (seasonal H3N2) | - | 1 (2.0%) | 1 (0.9%) | - | 2 (0.6%) |
| **Neuraminidase (NA)** | | | | | |
| A/California/07/2009 (H1N1/pdm09) | 29 (41.4%) | 27 (55.1%) | 57 (52.8%) | 77 (75.5%) | 190 (57.8%) |
| A/swine/Alberta/SD0217/2017 (H1N2 classical 1A.3.3.2) | - | - | - | - | - |
| A/swine/China/JG20/2019 (seasonal H3N2) | 6 (8.6%) | 7 (14.3%) | 11 (10.2%) | 22 (21.6%) | 46 (14.0%) |
| **Influenza B hemagglutinin (HA)** | | | | | |
| B/Austria/1359417/2021 | 2 (2.9%) | 1 (2.0%) | 5 (4.6%) | 7 (6.9%) | 15 (4.6%) |

**Supplementary Table 4.** Positive controls of influenza seropositive samples collected from human and ferrets.

| Sera | Subtype | Virus | Host |
|---|---|---|---|
| H1 | pdm09/H1N1 | A/Singapore/P019/2009 | Human |
| H3 | H3N2 | A/Singapore/P020/2009 | Human |
| H3 | H3N2 | A/Singapore/P146/2009 | Human |
| H3 | H3N2 | A/Singapore/P198/2009 | Human |
| H5 | H5N1 | A/duck/Laos/3295/2006 | Ferret |
| H7 | H7N9 | A/Guangdong/17SF003/2016 | Ferret |
| H9 | H9N2 | A/Hong Kong/308/2014 | Ferret |
| Negative control | - | Zero-bleed | Ferret |
